# Characterization of nuclear mitochondrial insertions in the whole genomes of primates

**DOI:** 10.1101/2020.02.24.963504

**Authors:** Gargi Dayama, Weichen Zhou, Javier Prado-Martinez, Tomas Marques-Bonet, Ryan E. Mills

## Abstract

The transfer and integration of whole and partial mitochondrial genomes into the nuclear genomes of eukaryotes is an ongoing process that has facilitated the transfer of genes and contributed to the evolution of various cellular pathways. Many previous studies have explored the impact of these insertions, referred to as NumtS, but have focused primarily on older events that have become fixed and are therefore present in all individual genomes for a given species. We previously developed an approach to identify novel Numt polymorphisms from next generation sequence data and applied it to thousands of human genomes. Here, we extend this analysis to 79 individuals of other great ape species including chimpanzee, bonobo, gorilla, orang-utan and also an old world monkey, macaque. We show that recent Numt insertions are prevalent in each species though at different apparent rates, with chimpanzees exhibiting a significant increase in both polymorphic and fixed Numt sequences as compared to other great apes. We further assessed positional effects in each species in terms of evolutionary time and rate of insertion and identified putative hotspots on chromosome 5 for Numt integration, providing insight into both recent polymorphic and older fixed reference NumtS in great apes in comparison to human events.

## INTRODUCTION

Polymorphic Nuclear Mitochondrial Insertions (NumtS) are fragments of mitochondrial DNA (mtDNA) that have been transferred and integrated into the nuclear genome of an organism. NumtS vary in size from smaller fragments to full length mitochondrial integrations and can share varying degree of homology with the genome of their parent mitochondria, depending on the age and time of insertion. Once these pieces of mtDNA have inserted themselves, their mutation rate decreases by an order of magnitude to the background rate of the organisms nuclear genome, essentially fossilizing the fragment and providing a snapshot of the ancestral mitochondria from when it was inserted(1–3). This information can be very beneficial in phylogenetic analysis(4, 5), and indeed these inserted sequences have been identified in previous studies exploring their prevalence in humans(5–7), chimpanzees(5, 8, 9), and other organisms(5, 10). As with other forms of genetic variation, NumtS can be both fixed within all genomes of a species as well as polymorphic between individuals. While, the majority of prior studies have focused on older, fixed insertions, recent efforts have begun to explore the impact of segregating Numt alleles within human populations(2, 7).

It has been shown that all parts of the mtDNA are able to transfer into the nuclear genome, though relative rates for individual regions are still unknown(7). Prior studies have reported a deficit of NumtS from the hyper variable (HV) regions of mtDNA, specifically HV2 of the mitochondrial control region (MCR_F_)(1). In particular, a negative correlation was reported between the prevalence of NumtS and the proportion of the variability of the site, thus increasing the difficulty of detecting events in nuclear genome that have a slower rate of mutation(1). More recent work focusing specifically on Numt polymorphisms, however, reported a slight enrichment of the D-loop for the polymorphic events(7). Likewise, the patterns of insertional preferences within chromosomal DNA have not been well defined. Earlier work reported that younger insertions are more prevalent within intronic regions while older NumtS tend to be intergenic(11). More recent analyses have suggested that they tend to insert non-randomly, with a tendency to insert in regions significantly deficit of transposable elements(12, 13) or the observation that in humans they seem to have a local affinity towards low GC regions and A+T dinucleotides(7), while others reported no pattern or specific insertion sites(14). An abundance of human specific NumtS on chromosome Y was reported in one study, though these appear to have arisen through the duplication of existing NumtS and were likely not driven through new insertions(1).

Previous studies have also found discrepancies in the rate of insertion for NumtS across different species. For example, some groups have found a consistent rate of insertion throughout the evolution of great apes(1, 6, 15), whereas others reported variability between species(9, 16). Indeed, earlier observations reported a higher rather rate of insertions in gorillas(17) and then later in chimpanzees(12) as compared to humans. Additional research has reported a burst of insertion at the divergence of old-world and new-world monkeys(5, 15, 18, 19). Further, an interesting divergence in the evolution of fixed Numt insertion was also reported between primate and non-primate species(5). Overall, there is much ambiguity regarding the interspecies rate of insertion, copy number and length, and these discrepancies are likely due in part to the predominant use of older, species-specific NumtS as well as limited sample size in the prior analysis.

Here, we present a holistic analysis of NumtS using both older, fixed insertions as well as new polymorphic events detected by our recently developed methodology(7). We derive the prevalence and insertion rates of NumtS across species and explore positional effects and sequence context. We further *de novo* assemble a subset of Numt insertions and analyze their evolutionary distance from constructed ancestral mtDNA sequences for the various speciation events. This represents the first exploration and assessment of polymorphic events and their significance in the evolution of the primates.

## METHODS AND MATERIALS

### Data acquisition and processing

Whole genome, paired-end sequencing data were obtained in SRA format from the short read archive (https://www.ncbi.nlm.nih.gov/sra) for various non-human primates: Chimpanzee – *Pan troglodytes*; Bonobo – *Pan panicus*; Gorilla – *Gorilla Gorilla* and Orang-utan – *Pongo abelii* and *Pongo pgymaeus*, were obtained from the Great Ape Genomes Project(20) and Rhesus - *Macaca mulatta* data were obtained elsewhere as an outgroup(21). The SRA files (SRP018689, ERP002376) were converted to fastq format using the SRA Toolkit (version 2.3.5-2 released April 2014) and then subsequently aligned to its respective species specific reference and processed using BWA(22) and GATK(23) to generate alignment files in BAM format. The reference versions used were as follows; for chimpanzee ‘panTro4’, for bonobo ‘panPan1’ (assembled by(24)), for gorilla ‘gorGor4’, for orang-utan ‘ponAbe2’ and for rhesus monkey ‘rheMac3’, and comparisons were made to NumtS previously identified in humans using the GRCh37 version of the human reference genome.

### Numt identification and assembly

In order to reduce misalignments from variability in mtDNA sequences within individuals of each species, modified references were generated for each sample by concatenating individual mtDNA to its species-specific reference. Specifically, sequence reads initially aligning to the reference mtDNA for each individual were isolated and assembled into a single mtDNA contig using CAP3(25). These sample specific mtDNA sequences were then inserted into each respective reference sequence as an additional contig and the sequence reads were realigned and processed as described above. The updated BAM files were used for polymorphic Numt discovery using *dinumt(7)*. NumtS in each species specific reference genome were reanalyzed using the method previously described(26) to generate a consistent and updated set across organisms.

The aberrant and soft-clipped read pairs from the flanking region of Numt insertion sites were then extracted and combined from all the samples with the insertion. Where possible, the polymorphic NumtS were assembled into contigs using CAP3(25). These contigs were subsequently aligned against their respective reference mtDNA genome to annotate the specific Numt fragment.

### Orthogonal validation of NumtS using long-read sequences

To assess the accuracy of putative NumtS insertion sites obtained from the short-read data sets, we made use of using existing long-read whole genome sequences. Long-read sequencing data from Pacific Biosciences (PacBio) single molecule, real-time (SMRT) technology were obtained from NCBI under the project accession numbers PRJNA369439 (chimpanzee, orangutan) and PRJEB10880 (gorilla)(27, 28). We used the pbmm2 (https://github.com/PacificBiosciences/pbmm2) to align the PacBio raw reads of three primate genomes to their relative references. We customized an approach for identifying mobile element insertions, PALMER(29), to allow the discovery of NumtS in PacBio aligned data. We required at least 4 sub-reads at a given locus with NumtS related sequences and performed additional error correction (sequencing error less than 1%) using CANU (https://github.com/marbl/canu)(30) to correct PacBio sub-reads that support the insertions. These were subsequently re-aligned locally to the predicted insertion sites using pbmm2, followed by a second run of PALMER to better define the sequences and integration sites of the detected NumtS. We then examined the intersection between calls from PacBio and from standard short-reads with an extension of ±500bp for insertion sites.

### Frequency and characterization of insertion site across all the species

To investigate whether insert-size or coverage had an effect on the number of insertions we observed, we derived and compared the frequency of Numt insertions relative to each reference for every non-human primate sample. The defined Numt breakpoints (site of insertion in the genome relative to the reference) were then unified to the human reference (GRCh37) using the lift-over tool and chain files from UCSC Genome Browser(31) to compare the sites of insertion across all the primates including humans(7) for both reference and polymorphic events.

A similar strategy was used to assess the subset of assembled NumtS. Their aligned coordinates were unified to the human reference (GRCh37) as described above and a circos plot was generated to display integration preferences for mtDNA fragments into each species. The human reference mtDNA (GRCh37) was binned into genes and the average rate of insertion for each bin was calculated across all species.

### Characterization and rate of insertions for NumtS

To investigate species-specific differences in insertional characteristics, we compared the fragment length and GC content for all the assembled polymorphic and reference NumtS (extracted from reference genome of each species). In addition, the rate of insertion was estimated for assembled polymorphic NumtS using the method described previously(7). The ancestral mitochondrial sequences for each primate genus were obtained from ENSEMBL Compara release 77 based on 8 primates EPO (Enredo-Pecan-Ortheus) pipeline(32).

### Enrichment analysis

To test whether there were positions within each species that were predisposed for Numt insertions, we conducted a permutation analysis by generating 1000 sets of random insertion coordinates based on the list of polymorphic and reference Numt fragment sizes. The frequency of complete gene insertion were calculated for all the MT genes for each species. Since some of the primate species were lacking the MT gene annotation, we defined the genes using human mtDNA reference (GRCh37) for each primate species through the lift-over tool and chain files from UCSC Genome Browser(31). The Z-score was calculated for each gene to determine trends of MT gene enrichment or depletion across all human and non-human primates as follows

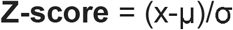

where x is the true values of total gene insertions, μ is the mean of permutation count and σ is the standard deviation of the permutation count.

### Analysis of putative hotspots of Numt insertion

The entire genome was divided into 25 Mbp bins and the frequency for Numt insertions was calculated and compared across the homologous positions in each species to determine any hotspots above a permutation threshold that defines 5% FDR (false discovery rate) as follows

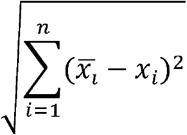

Where is the mean frequency of NumtS across all the bins for a given species *i* and is the frequency of NumtS in each bin, equated across number of species. 25 Mbp was determined to be a lower bound for window size given the sparsity of reference and polymorphic NumtS in each species.

## RESULTS

### Discovery and frequency of NumtS across species

We identified a total of 409 polymorphic NumtS in 79 samples across 5 different non-human primate species (**Additional file 1**). Chimpanzee was observed to encompass the largest number of polymorphic Numt insertions with a total of 187 events discovered in 24 samples across four different subgroups at an observed rate of 20.8 average per sample. *Pan troglodytes troglodytes*, *P. troglodytes ellioti*, *P. troglodytes schweinfurthii* exhibited an average of 22 events per sample, though *P. troglodytes versus* showed a much lower rate of 10.2. Macaque had the highest overall average per sample rate of 21.2 insertions and a total of 56 unique events called in 5 samples, while organ-utans showed the lowest at 12.9 average per sample, with 43 total unique events in 10 samples from two different species though with differing rates between *Pongo Pygmaeus* (17.2 avg/sample) and *Pongo abelii* (8.6 avg/sample). The bonobo individuals in aggregate had 67 total insertions with an average of 15.46 per sample and 27 samples of gorilla with total of 55 Numt calls belonging to 2 different species and 3 different sub-species had an overall 14.6 ave/sample. Both *Gorilla gorilla gorilla* and *G. gorilla diehli* had approximately 15 events per sample, but the sub-species *G. beringei graueri* only had 2.5 ave/sam similar to human polymorphic events (**Table 1**). We are also reporting a total of 4274 reference (fixed) NumtS in great apes and macaque (chimpanzee - 974, bonobo - 809, gorilla- 657, orang-utan - 963, and macaque - 871) (**Additional file 2**).

**Table 1:**
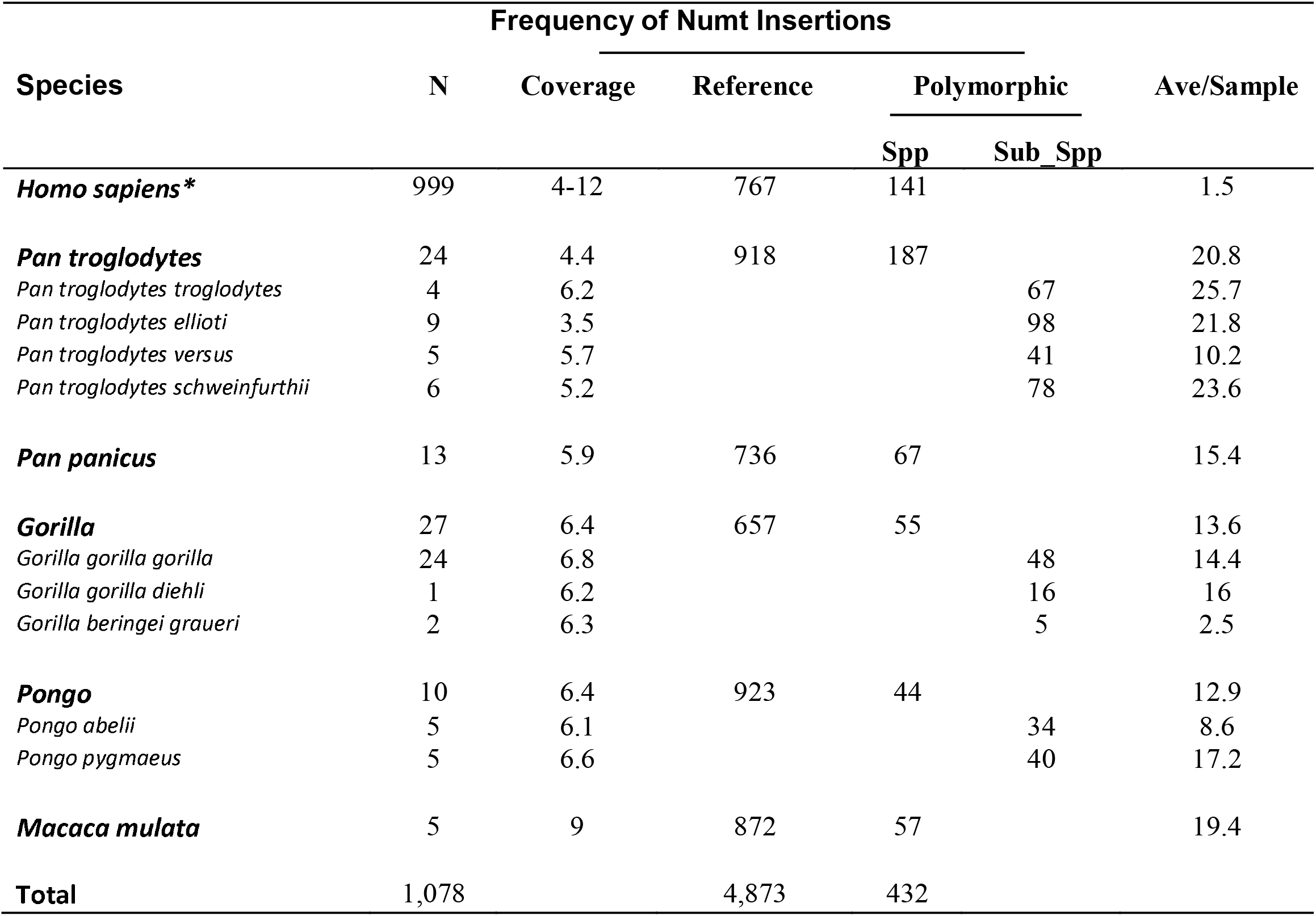
Frequency of NumtS. The frequency of Numt insertions in reference genomes and Numt insertions relative to the respective reference, discovered in several (N=number of samples) samples of 6 different groups (Human, Chimpanzee, Bonobo, Gorilla, Orang-utan and Macaque). Chimpanzee and gorilla are further divided into subspecies. Therefore, the total polymorphic NumtS for the species are listed under “spp” where as the frequency for subspecies are listed under “sub_spp”. Average coverage for each group is noted in the third column. The last column has the average Numt insertions in each group.

We next assessed the frequency of NumtS that were unique to each sample and observed a similar trend in chimpanzee, bonobo and macaque with a higher rate of sample specific, relatively recent insertions compared to other species. In contrast, most insertions in gorilla and orang-utan were shared among other members of the species (**Figure 1**). We also discovered several events that were found in the majority of samples suggesting older, fixed insertions. We observed an overall higher frequency of polymorphic insertions in all the chimpanzee sub-species, with the exception of *P. troglodytes versus* which exhibited a frequency similar to the other primates. Furthermore, we did not observe any effect of insert-size or coverage on the number of insertions (**Additional file 3: Figure S1).**

**Figure 1:**
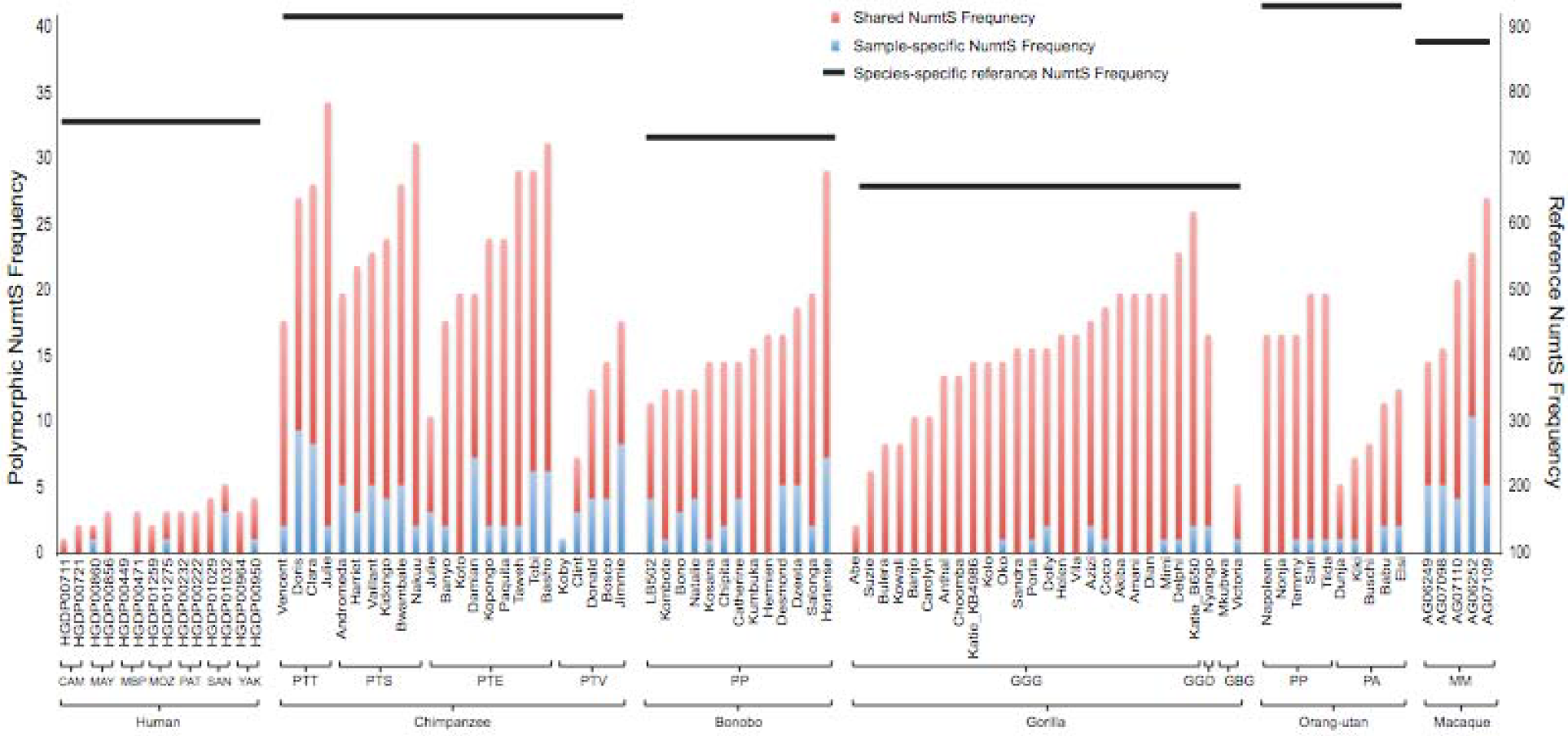
Shared and sample specific Numt frequency. Each bar is representing the frequency of Numt insertion in that sample. The red part of the bar is to show the number of events that are shared between samples within that species, where as the blue bar is for the Numt insertions unique to that sample. A three letter code is used for grouping the subspecies (in case of chimpanzee and gorilla) and population (for humans), where as a two letter code is used for denoting the species (for bonobo, orang-utan and macaque).

### Spatial organization of Numt insertions into nuclear primate genomes

We examined the chromosomal locations of Numt integration across homologous regions of each species in order to explore whether there were shared insertional preferences. We found no indication that any specific regions harbored an extreme enrichment or depletion of either NumtS present in the respective references of each species or polymorphic events (**Additional file 3: Figure S2A and S2B**), with the exception of chromosome Y for which no polymorphic insertions were detected in the species in which this chromosome was available for analysis (humans, chimpanzees and orang-utans). There were also several gaps on some chromosomes in different species that may be due to assembly differences, particularly around the centromeric regions, resulting in a dearth of Numt detection.

### De novo assembly of Numt insertions

We next sought to derive the underlying sequences of the Numt insertions. In contrast to our earlier work with human cell lines, the underlying primate samples from which the whole genome sequences were derived were either unavailable or very sparse and thus prohibit the direct molecular characterization of these events. We thus applied an assembly strategy (see Methods and **Figure 2**) around each putative insertion site and were able to construct assembled contigs for 54% of our predictions across each species. We assessed the accuracy of our assembled sequences through comparison with high quality NumtS we previously sequenced and observed a very high concordance at concordant sites (99.9% average identity). Using this approach, we were able to recapitulate the underlying insertion sequences for 222 NumtS across all species (**Additional files 4-8)**. The majority of NumtS that we were able to assemble were found in multiple samples. Longer/full-length NumtS were more difficult to assemble due to the circular nature of mtDNA as well as the limitations of using short reads, though we were able to identify the possible length and region of MT for long insertions based on the partial assembly of each breakpoint.

**Figure 2:**
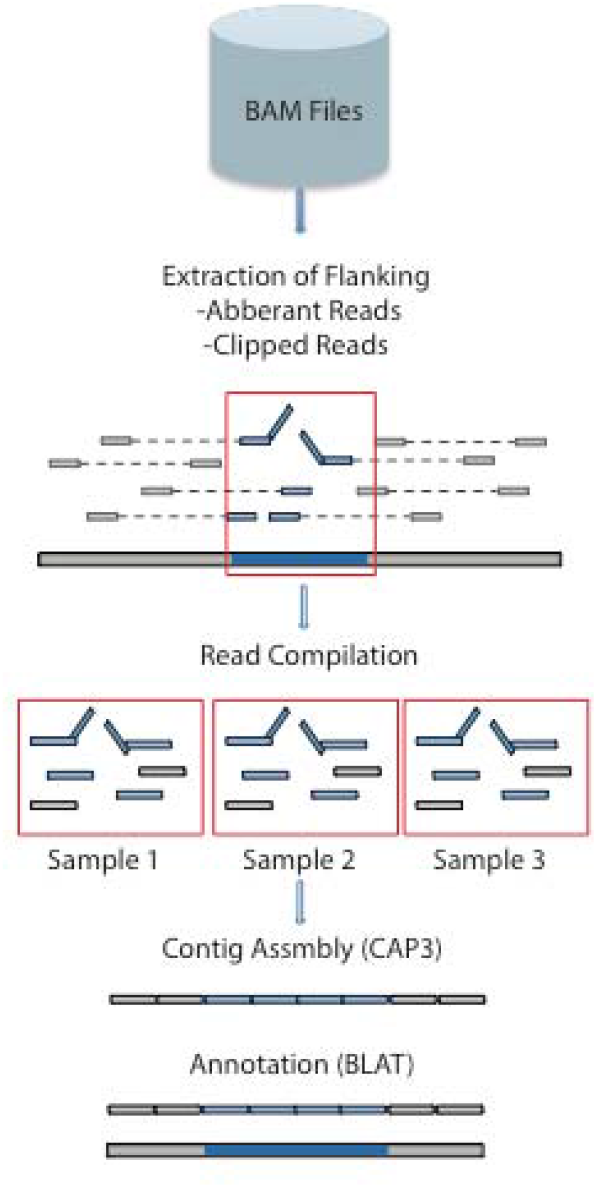
Pipeline for Numt assembly. The graphical representation is divided into three main parts of the Numt assembly pipeline. A) Utilizing BAM files to extract the flanking reads and split reads around the breakpoints of Numt insertion from all the samples that they were discovered in. B) These reads are then combined into one file to assemble into contigs using CAP3. C) The contigs are then annotated using UCSC BLAT feature.

### Validation of Numt calls using long-read sequencing

PacBio long-read sequencing results in longer (10 Kbp or higher on average) contiguous DNA sequences than standard short-read sequencing technologies and can identify full-length insertions that fall completely within an individually sequenced molecule (Chaisson et al. 2017; Sedlazeck et al. 2018). We made use of three primate genomes (one Chimpanzee, one Gorilla, one Orangutan) sequenced to >65X by PacBio to provide orthogonal validation for our call sets of NumtS(27). Although these sequences were obtained from different individual samples and were not present in the short-read cohort we analyzed, it was still likely that we would identify some species-specific polymorphisms that were shared between the sets that could be used as a basis for comparison. Indeed, we observed orthogonal evidence for 7 NumtS in Chimpanzee, 4 NumtS in Orangutan, and 1 in Gorilla (**Additional file 9**) from PacBio data. The PacBio long reads also provided augmented information for these calls, including the specific sequence and segment coordinates within the mtDNA as well as insertion orientation (**Additional file 9**). In addition to verifying the insertion loci in the nuclear genome, we further compared our assembled NumtS to the sequences present in the respective PacBio long reads and found a 99.5% average identity concordance between the contigs. Together, the results showed a high consistency between calls by PacBio data and our assembled Numt insertions by standard short-read technology with regard to insertion loci, segment coordinates within the mtDNA and the insertion sequences, suggesting we obtained a high confidence set of polymorphic assembled Numt insertions.

### Characterization of Numt insertional preferences in the nuclear genome

We next investigated whether there was any enrichment for polymorphic or reference NumtS within or across the different species. We segmented the human genome (GRCh37) into bins of 25 Mbps and compared the frequency of Numt insertion (lifted over to the human reference) for each bin across all the groups (human, nonhuman primates and macaques) against a random background. Eight hotspots with a frequency greater than the threshold value (permutation threshold that defines 5% FDR) were observed for reference NumtS, one of which on chromosome 5 was also identified as the sole hotspot for polymorphic insertions (**Additional file 3: Figure S3**). We further explored this latter region (chr5:75,000,000-100,000,000) to determine whether this was specific to an individual species are common across primates. We observed that the majority of these insertions were primarily from chimpanzee, bonobo and gorilla, while human, orang-utan and macaque were lacking any insertions in this region for polymorphic events. However, we did observe an overall high abundance of fixed reference events in all six species, with orang-utans exhibiting the largest number of insertions and humans the least.

### Characterization and enrichment of NumtS

We previously reported that polymorphic NumtS in humans exhibited a higher GC% than their parent mitochondrial genome(7), and thus we sought to determine whether this observation extended to non-human primates as well. We calculated the GC% for the subset of NumtS we were able to assemble for each species and unlike humans, observed no significant change in any of the polymorphic insertions compared to their parent mtDNA with the exception of those present in chimpanzee (**Figure 3**). However, the GC% for all polymorphic events differed between species, with those within macaque, gorilla and bonobo exhibiting a lower GC% in contrast to the higher GC% observed in human, chimpanzee, and orang-utans (**Additional file 3: Figure S4)**. While a similar trend in average GC% was also observed for reference mtDNA in all species, there was a gap observed only between GC% of polymorphic events and their respective reference mtDNA for chimpanzee and human. This suggests that while polymorphic events mimics the GC% of their parent mtDNA in all species, chimpanzees and humans show a higher GC% for recently inserted polymorphics NumtS compared to their parent mtDNA. We further generated a phylogenetic tree using the reference mtDNA for each species and their respective ancestral mtDNA to explore ancestral relationships (**Additional file 3: Figure S5).** Additionally, we also found that the average GC% for macaque, orang-utan and human ancestral mtDNA was lower compared to GC% of polymorphic NumtS and reference mtDNA, whereas it was similar or higher for the rest of the species (**Figure 3**).

**Figure 3:**
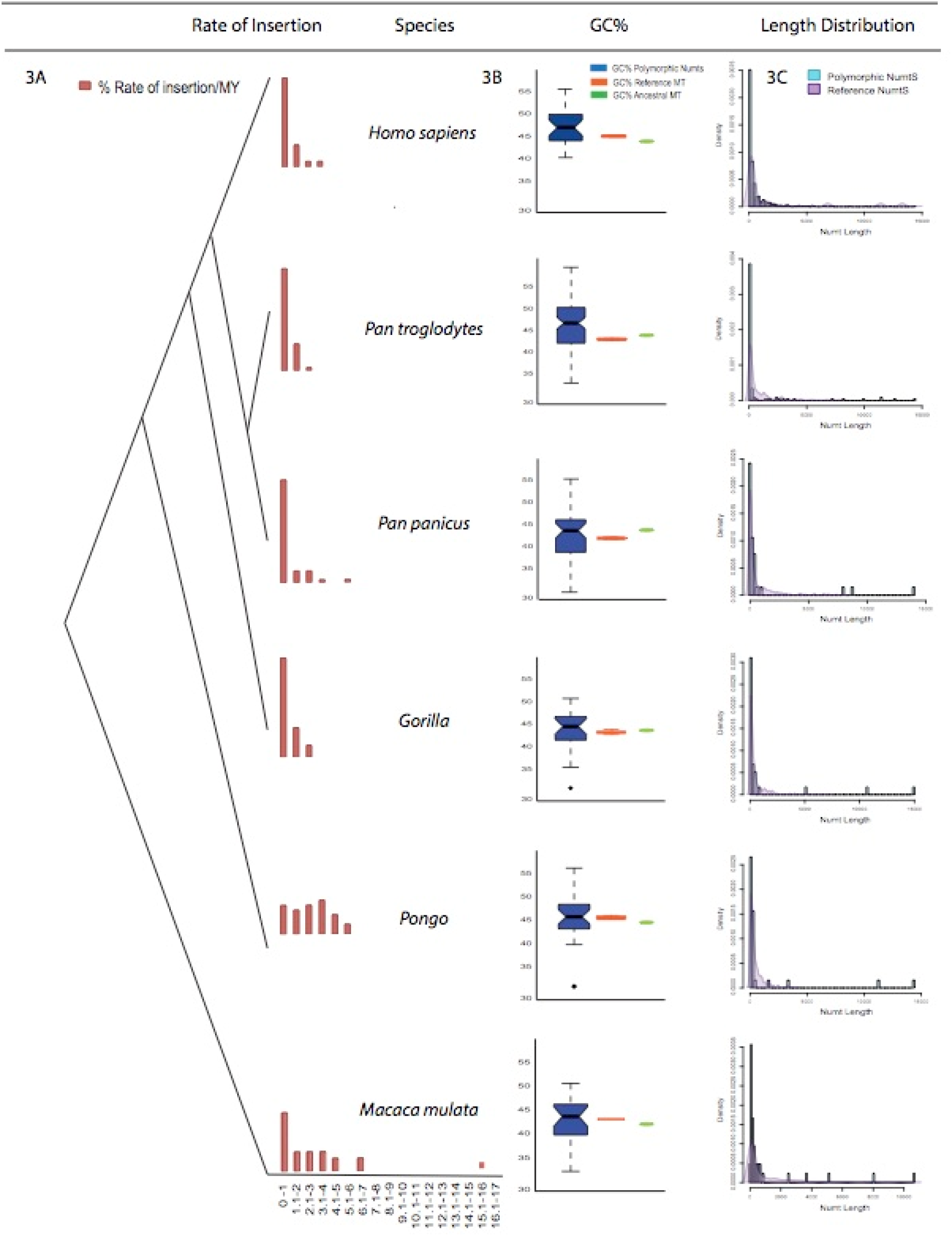
Rate of insertion, length of NumtS and GC content. A) Rate of Numt insertion represented by the red bars divided per MYs for all 6 groups. B) %GC content in assembled polymorphic Numts compared (represented by blue notched box plot) with %GC content of reference mtDNA (orange bar) and ancestral mtDNA (green bar). C) Length of assembled polymorphic NumtS represented by the blue bars compared to length of reference NumtS represented by overlain purple density plot.

We assessed whether there were length differences between our polymorphic NumtS and those present in their respective reference sequences (**Figure 3**). The majority (~95%) of the assembled polymorphic insertions were smaller than 1Kb in length, which is expected given the limitations of short-read assembly and is consistent with our previous results(7). Even though we were not able to fully assemble these long insertions, we were able to partially assemble contigs that crossed over the breakpoints precisely and which also matched our predicted mtDNA coordinates, enabling us to infer that ~5% of the events were likely full length. In contrast, reference insertions across all species were up to 8Kbp in size, with 3 (>8Kbp) outliers in only humans.

We examined whether there were any insertional preferences for any particular region of mtDNA, with a focus on Numt fragments containing complete genes or the D-loop (**Figure 4A and 4B**). We compared detected NumtS in the reference sequences of each species to random background models of insertions by converting gene annotations for mitochondrial genes for each species relative to human mtDNA using the liftOver tool (see Methods) and did not observe preferential bias towards any region or genes from the mtDNA relative to the human reference mtDNA sequence. We next examined polymorphic and reference NumtS identified individually at the species level and observed several regions of mtDNA that exhibited enrichment or depletion across multiple genes (**Additional file 3: Figure S6-S11**). We assessed these regions across all species to identify any selective biases towards particular MT genes and did not observe any pan-species MT gene bias for reference NumtS (**Figure 5A).** However, for polymorphic events we found an enrichment of genes located towards the center of mtDNA and depletion closer to the D-loop (**Figure 5B**). This was consistent across all species except bonobo, which interestingly exhibited the opposite showing a depletion of genes towards the center of mtDNA and enrichment of D-loop containing fragments.

**Figure 4:**
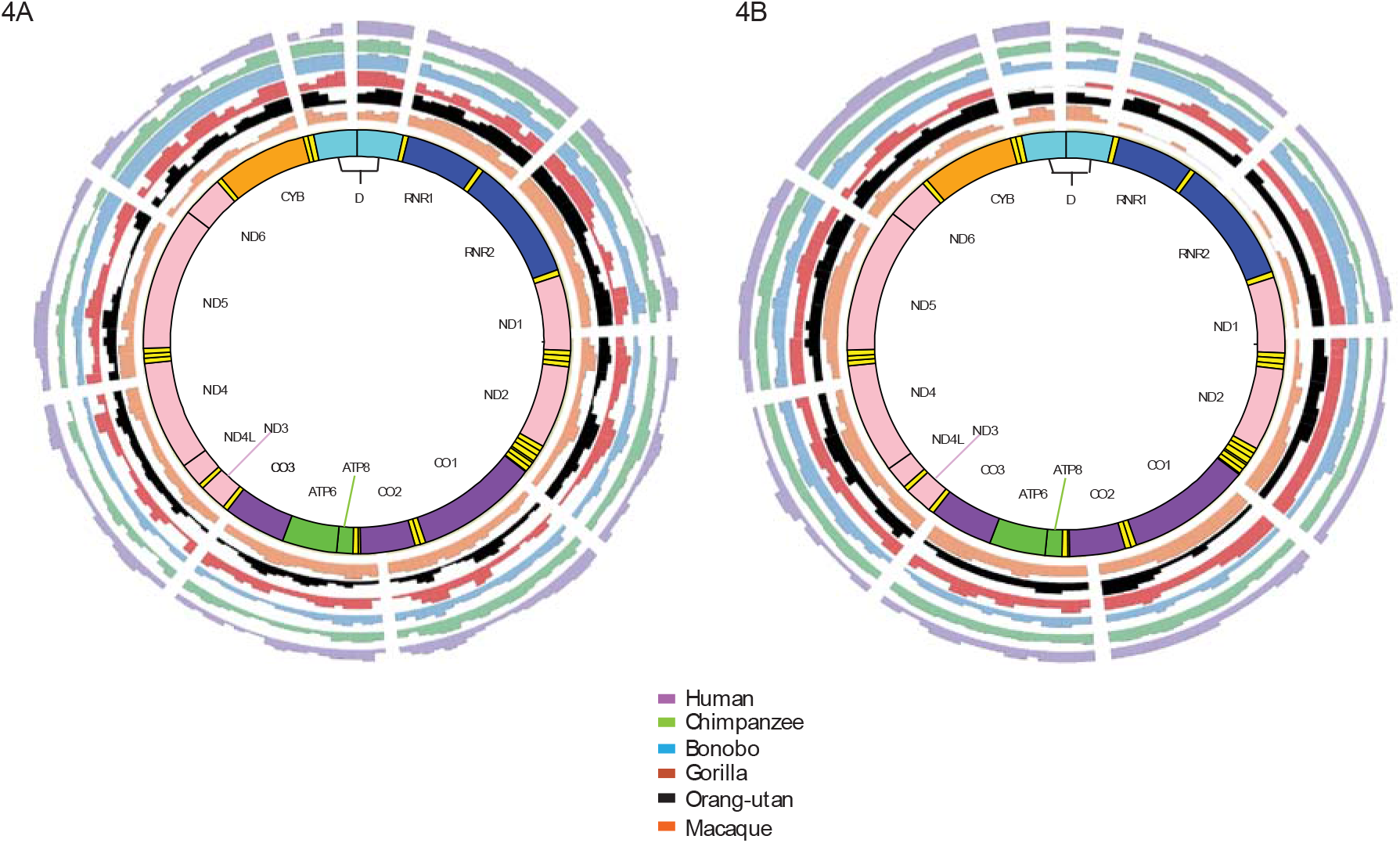
Spatial representation of species-specific insertions relative to the human reference genome. The breakpoints for all the species are lifted over to human reference (GRCh37/hg19). A) The normalized frequency of each mitochondrial gene across all the reference Numt fragments. Unique color is assigned for each genus, purple for humans, green for chimpanzee, blue for bonobo, red for gorilla, black for orang-utan and orange is for macaque. Using the same color scheme figure (B) was generated to illustrate the frequency of polymorphic Numts.

**Figure 5:**
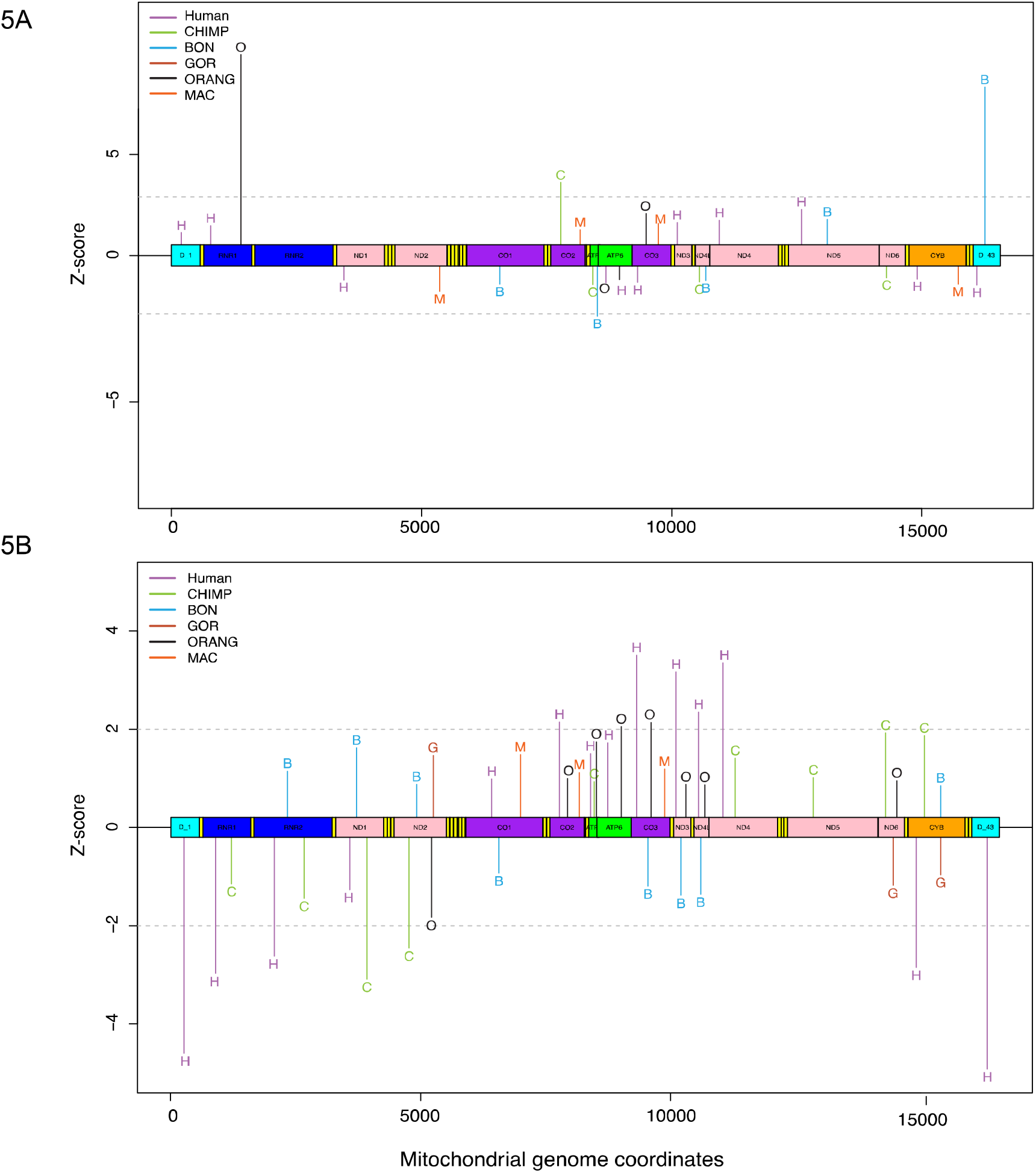
Mitochondrial gene enrichment or depletion across species. A) Linear mtDNA representing the enriched and depleted (Z-score >= 1 & <= −1) major genes (genes coding for tRNA excluded) inserted as polymorphic NumtS in all 6 species. Unique color and alphabet is assigned for each genus, purple ‘H’ for humans, green ‘C’ for chimpanzee, blue ‘B’ for bonobo, red ‘G’ for gorilla, black ‘O’ for orang-utan and orange ‘M’ is for macaque. B) Using the same color scheme figure (B) was generated to illustrate the significant Z-scores for polymorphic Numts.

### Rate of insertion

Once a piece of mtDNA inserts itself into the nuclear genome, it typically follows the mutational rate of its host DNA that is a magnitude lower than the mtDNA, giving a snapshot of when the insertion occurred(1–3). Based on this, we calculated the rate of insertion per million years normalized against the evolutionary time of divergence. We observed that most polymorphic insertions occurred in the past million years for gorilla, bonobo, chimpanzee and macaque, similar to the humans. Orang-utan was the only group that showed a different rate of insertion, where the insertions were spread evenly over the past 4 to 5 million years (**Figure 3**). Interestingly, we also found a Numt that inserted ~15 MYA in macaque. It is possible however that these insertions are fixed in the population but are not a part of the reference due to weak characterization and gaps in some of these primate species.

## DISCUSSION

Advances in sequencing technologies have made it possible to conduct studies at a finer scale, allowing the exploration of previously understudied genomic variation in populations across multiple species. Here, we have described one of the first forays into the investigation of polymorphic Nuclear Mitochondrial Insertions (NumtS) in multiple samples from primates and an old world monkey. We report a shift in the frequency of polymorphic Numt insertions with a successive increase from orang-utans to chimpanzees, followed by a decrease in humans by an order of magnitude. The highest rate of insertion was observed in macaques which diverged from humans approximately 25 MYA ago. This suggests that at some point after macaque-primate divergence there was a considerable drop in the rate of insertion as seen in orang-utans. Indeed, a burst of Numt insertions in the reference genome (and thus, primarily fixed in the species) has been previously reported(15) around the same time period as the divergence of old world monkeys, which is consistent with their presence as an outgroup in our analysis. Although we can posit no specific cause for the change in the rates of insertions, we can proffer several speculations. First, this might be due to Mitochondria’s rate of disintegration caused by factors such as oxidative phosphorylation(33) unique to each species. Alternatively, the prevalent mechanism for Numt insertions is understood to be implicated with the repair of double stranded breaks (DSB)(8), however there may be other mechanisms yet to be discovered that are specific to a given species.

We also observed an absence of polymorphic Numt insertions on Y chromosome across all species. This is in part due to the bonobo, gorilla, and macaque reference genomes omitting the Y chromosome, however we also did not observe any polymorphic insertions in orang-utan, chimpanzee or even humans whose references are more complete. This suggests some selective pressure against these insertions on Y chromosome; if the insertions were to occur in the gamete stage, when only the ovum contains mtDNA and not the sperm, the absence of any insertions on the Y chromosome might be explained. This hypothesis is further supported by the findings shown by(18) that all the recent reference NumtS on the Y chromosome were due to the result of duplication rather than novel insertions.

We observed a higher GC% of polymorphic NumtS in chimpanzee and humans compared to their respective parent mtDNA, which might be due to two reasons; 1) If there is an insertional bias towards mtDNA fragments with higher GC%, to balance the low GC content of the region where the insertion occurs on nuDNA or 2) the mtDNA fragment with lower GC disintegrating faster than the fragments that have a higher GC, thus increasing the chances of these fragments being selected for repairing the non-homologous breaks.

We further report an enrichment of both reference and polymorphic Numt insertions on chromosome 5. While for reference events the hotspot was derived by substantial number of insertions across all the species (especially orang-utan with 50 insertions), polymorphic events were mainly derived by gorilla and bonobo. We did observe that human reference (GRCh37) has a slightly lower average GC% at 34% for 100bp flanks on either side of Numt breakpoints in this region. Further,(34) examined structural variations and transposable elements by dividing the genome into 10Kbp bins and reported several hotspots in this 25mb region for human (9), chimpanzee (6) and orang-utan (6). Interestingly, they did not find any hotspots in rhesus. So far the only known mechanism for Numt insertions are through repair of DSBs, but exploring this phenomenon, could potentially lead to other mechanistic explanations for Numt insertions and for their continued occurrence.

It should be noted that the above mentioned observations might also possibly be because some of the primate references are not well defined and we might be missing either newer NumtS relative to the reference or calling events that are fixed in the species but missing in the reference. We also note that there is likely an ascertainment bias however due to the preferential assembly of smaller NumtS events from the shortread sequence data available.

Our study does have some limitations, such as the lack of direct molecular validation due to the unavailability of most of the samples we analyzed. However, we previously established the high fidelity of our NumtS detection method in humans(7) and expect a similar accuracy for the non-human primates assessed here. We also made use of long-read PacBio sequences which, while limited, had very good concordance with both our predicted insertions as well as their assembled sequences. This indicates that our *de novo* assembly approach can serve as a valuable tool for when exploring such insertions in nuclear genomes for datasets where further molecular characterization of samples is not possible.

## ABBREVIATIONS

Numt: Nuclear insertions of mitochondrial origin
mtDNA: mitochondrial DNA
nuDNA: nuclear DNA
DSB: double stranded breaks
DBSR: double stranded break repair
MYA: million years ago
FDR: false discovery rate.

## ACKNOWLEDGEMENTS

We would like to thank Jeffrey M. Kidd for his insightful advice on the analysis and also Christian Brion for help with figure construction.

