## Additional file 3 for "Characterization of nuclear mitochondrial insertions in the whole genomes of primates"

### Supplemental Figures:

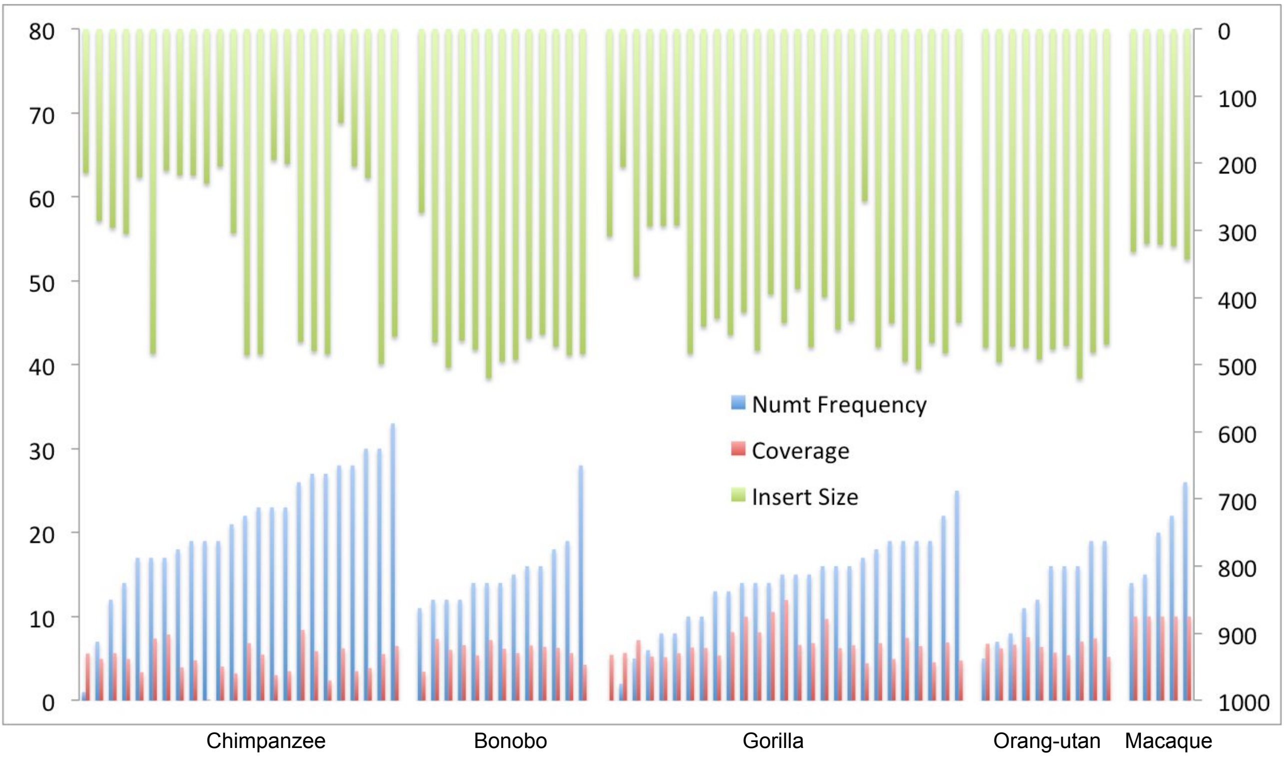


**Fig S1. Numt frequency.** The frequency of polymorphic Numt insertions (blue bars) relative to the sample coverage (red bars) and average insert size in bp (green bars) discovered in samples of 5 different groups (Chimpanzee, Bonobo, Gorilla, Orang-utan and Macaque).

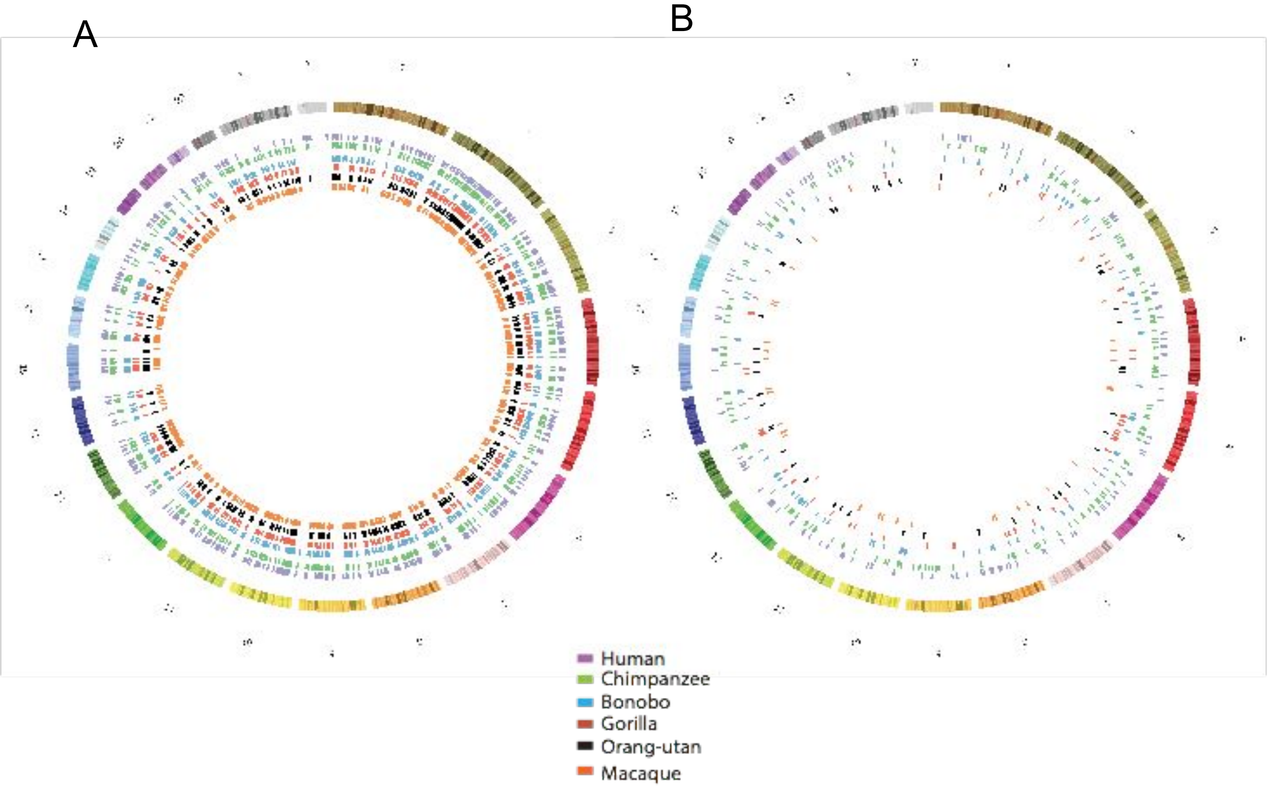


**Fig S2. Spatial representation of species-specific insertions relative to the human reference genome.** The breakpoints for all the species are lifted over to human reference (GRCh37/hg19). A) Position of Numt insertions for reference events. Unique color is assigned for each genus, purple for humans, green for chimpanzee, blue for bonobo, red for gorilla, black for orang-utan and orange is for macaque. Using the same color scheme figure B were generated to illustrate the Numt insertions for polymorphic events.

A

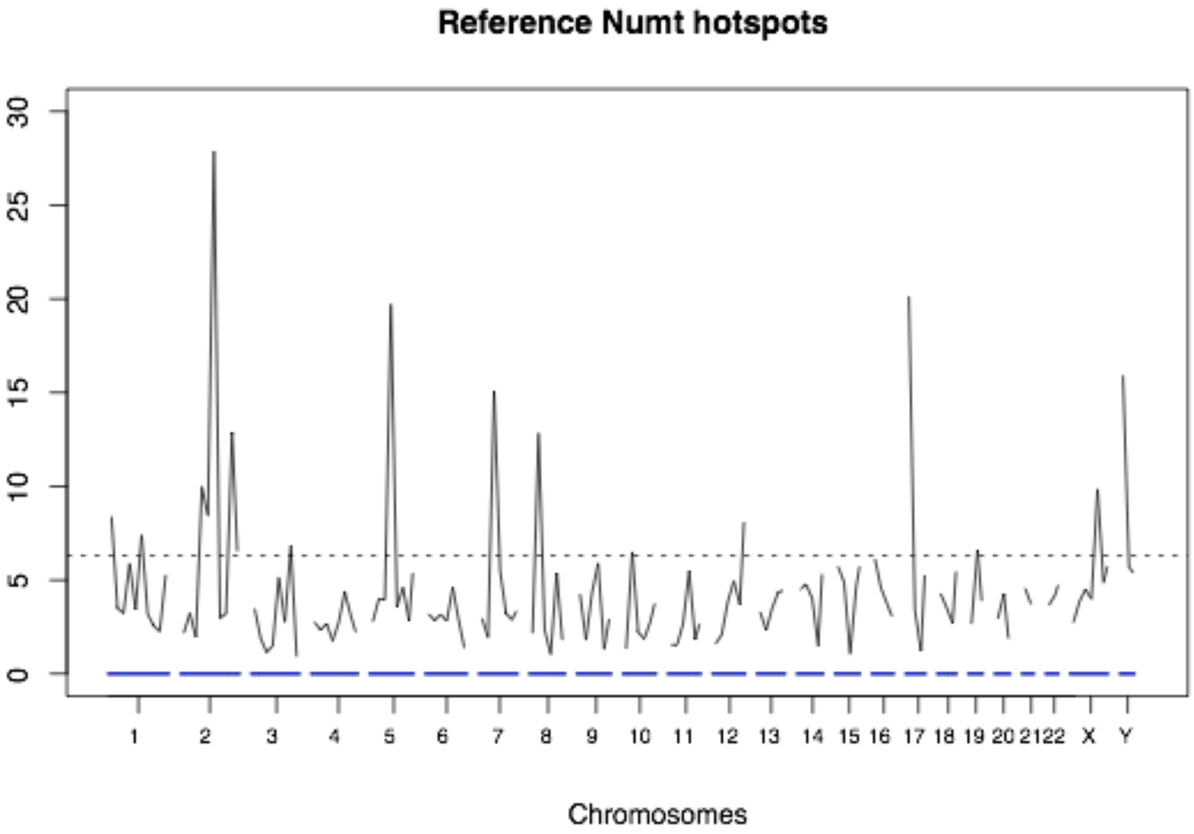


B

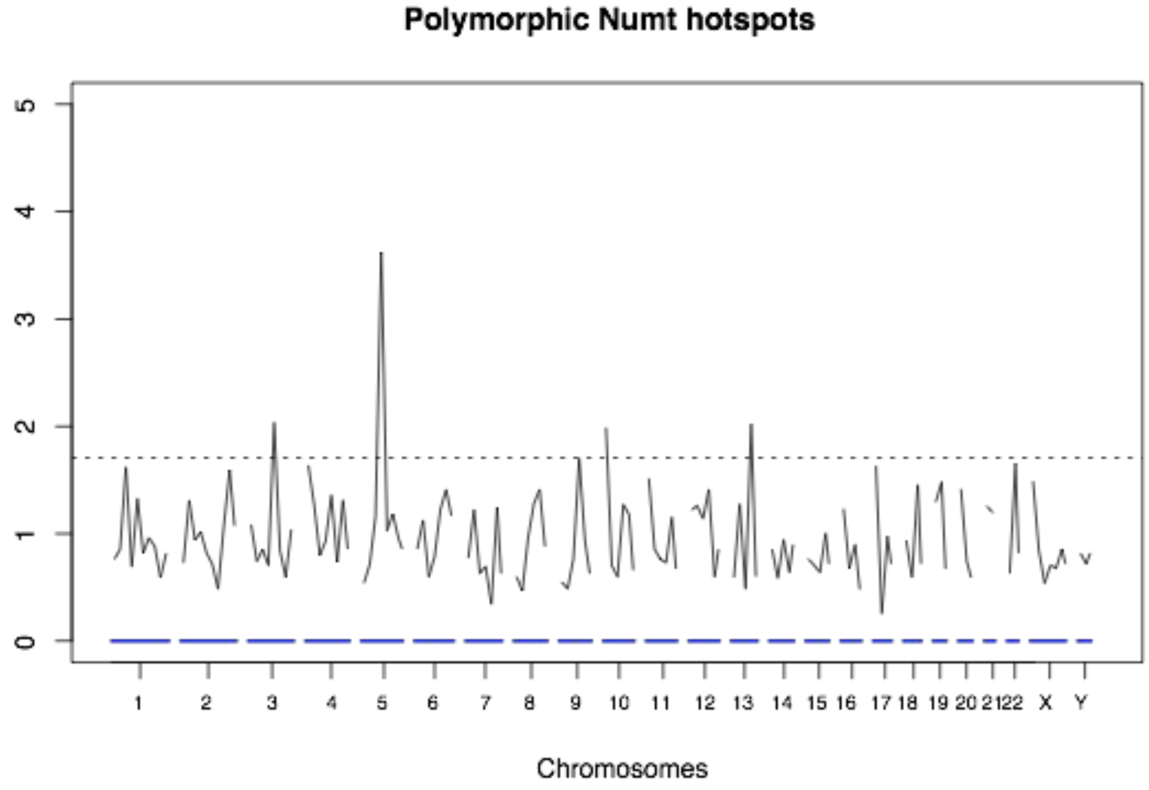


**Fig S3. Hotspot analysis**. Frequency of A) reference and B) polymorphic Numt events along the genome, compared across the homologous positions of the 6 species (see methods for hotspot frequency calculation). Each chromosome are divided into 25mbp bins. The dotted line is permutation threshold that defines 5% FDR (false discovery rate) of significant hotspots.

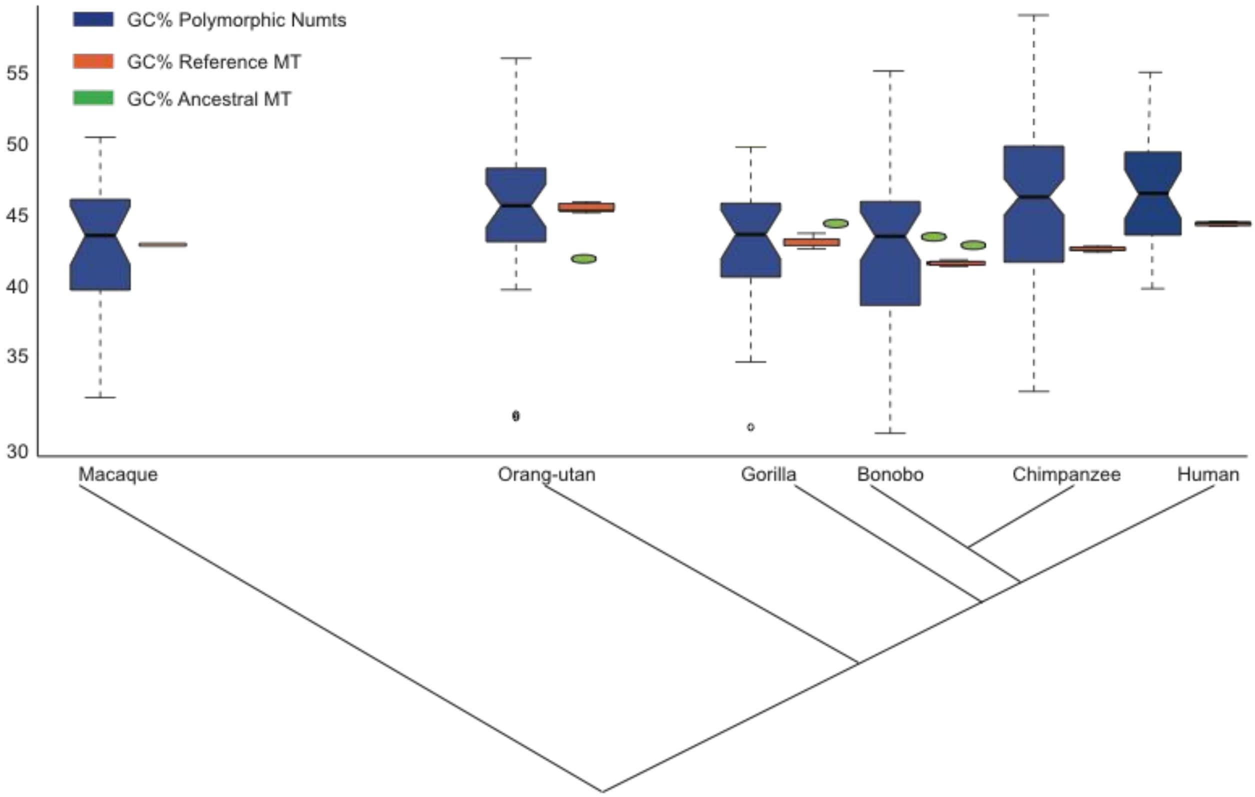


**Fig S4. %GC for polymorphic NumtS, reference mtDNA and ancestral mtDNA**. Comparing the GC% of polymorphic NumtS (blue) from each species to the average GC% of their respective reference mtDNA (orange) and respective ancestral mtDNA (green).

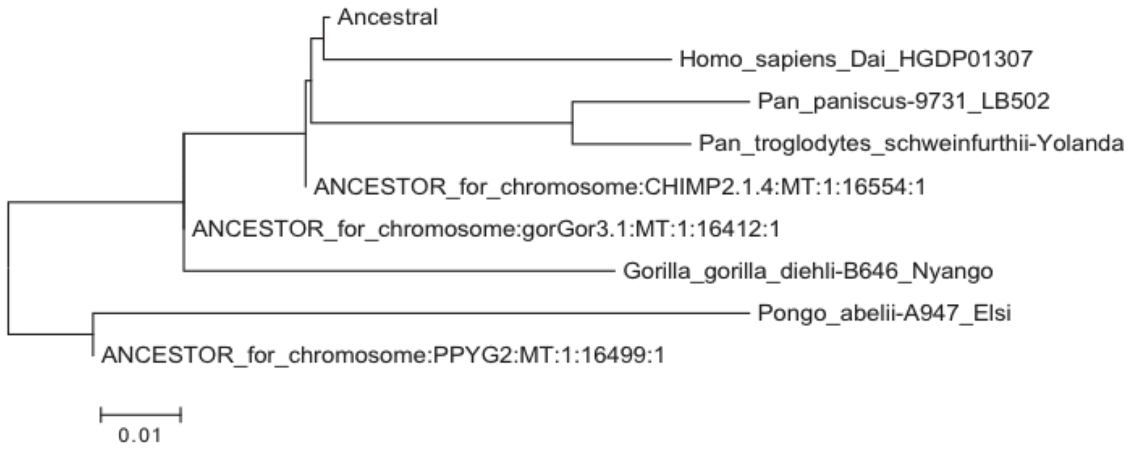


**Fig S5. Phylogenetic tree**. Comparison of ancestral and reference mtDNA to validate the link at each node of speciation.

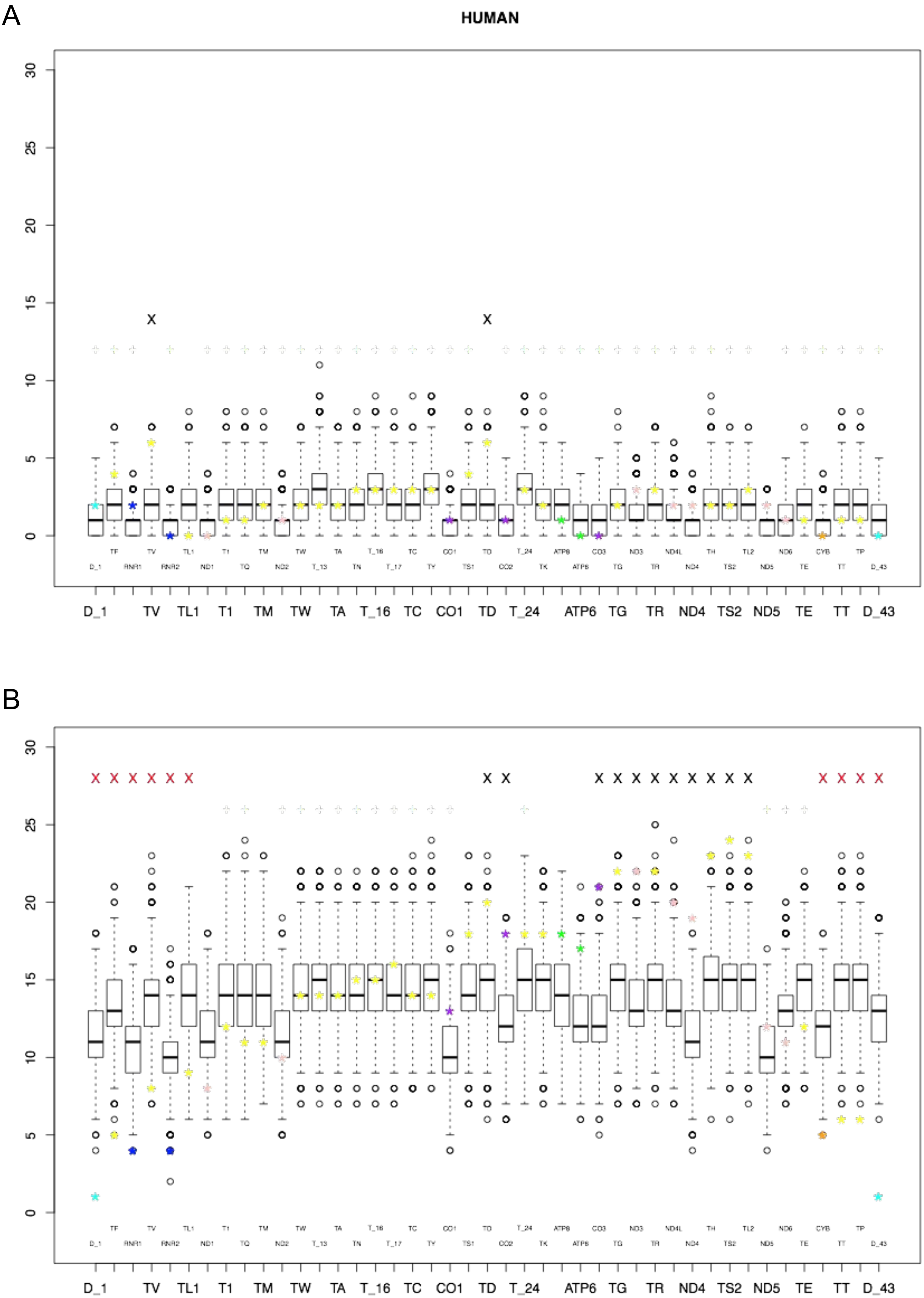


**Fig S6. Enrichment analysis for complete mitochondrial genes integrated into the human genome, relative to human mtDNA sequence**. Observations are indicated by asterisks and are relative to 1000 permutations of random mtDNA fragments of the same size (box plots). Significantly enriched or depleted genes (p-value <= 0.05) are denoted with a black or red X, respectively. Results are shown for (A) reference-based and (B) polymorphic NumtS.

A

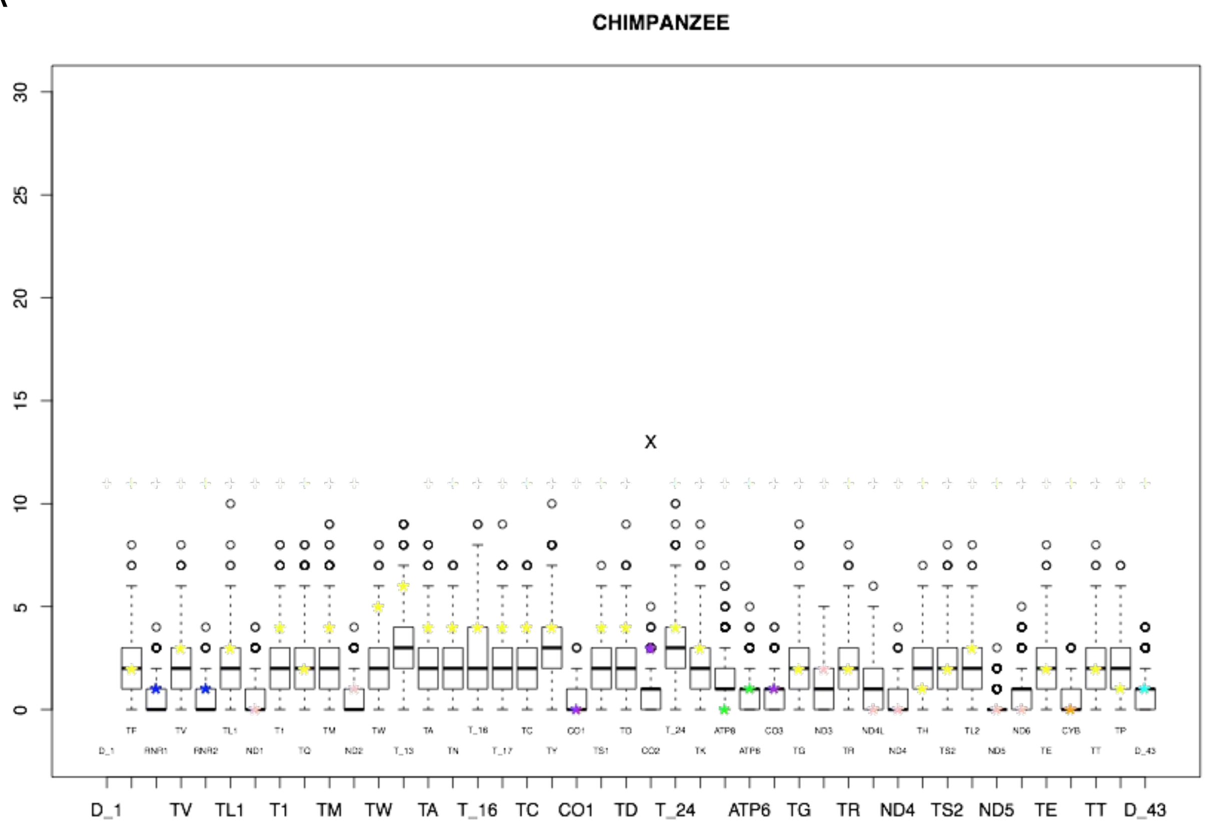


B

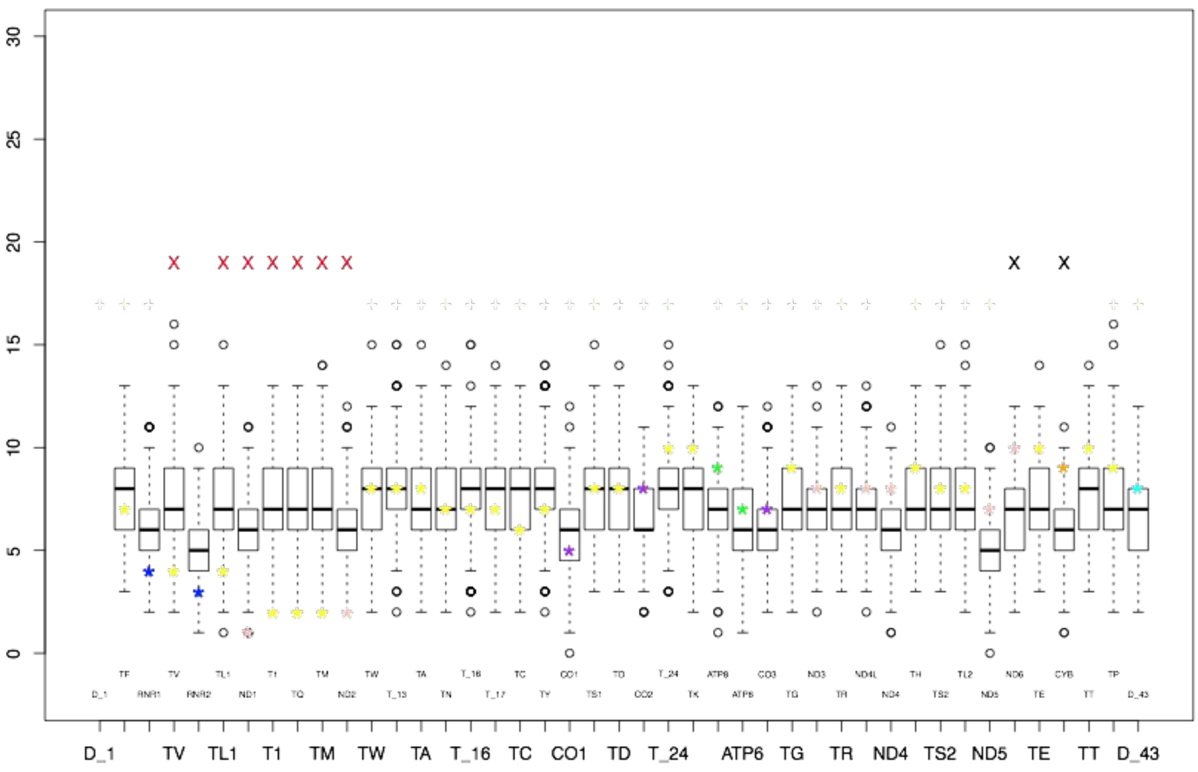


**Fig S7. Enrichment analysis for complete mitochondrial genes integrated into the chimpanzee genome, relative to human mtDNA sequence.** Data represented as described in Fig S3 for (A) reference-based and (B) polymorphic NumtS.

A

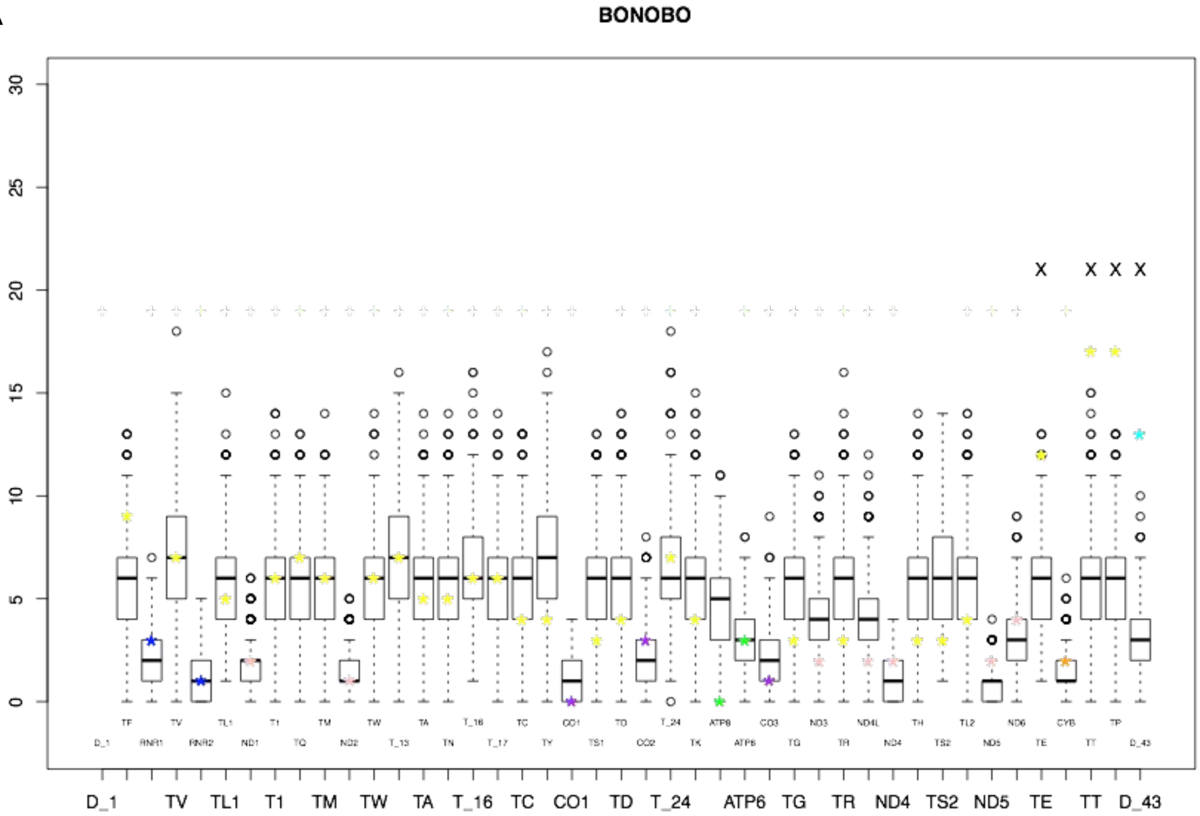


B

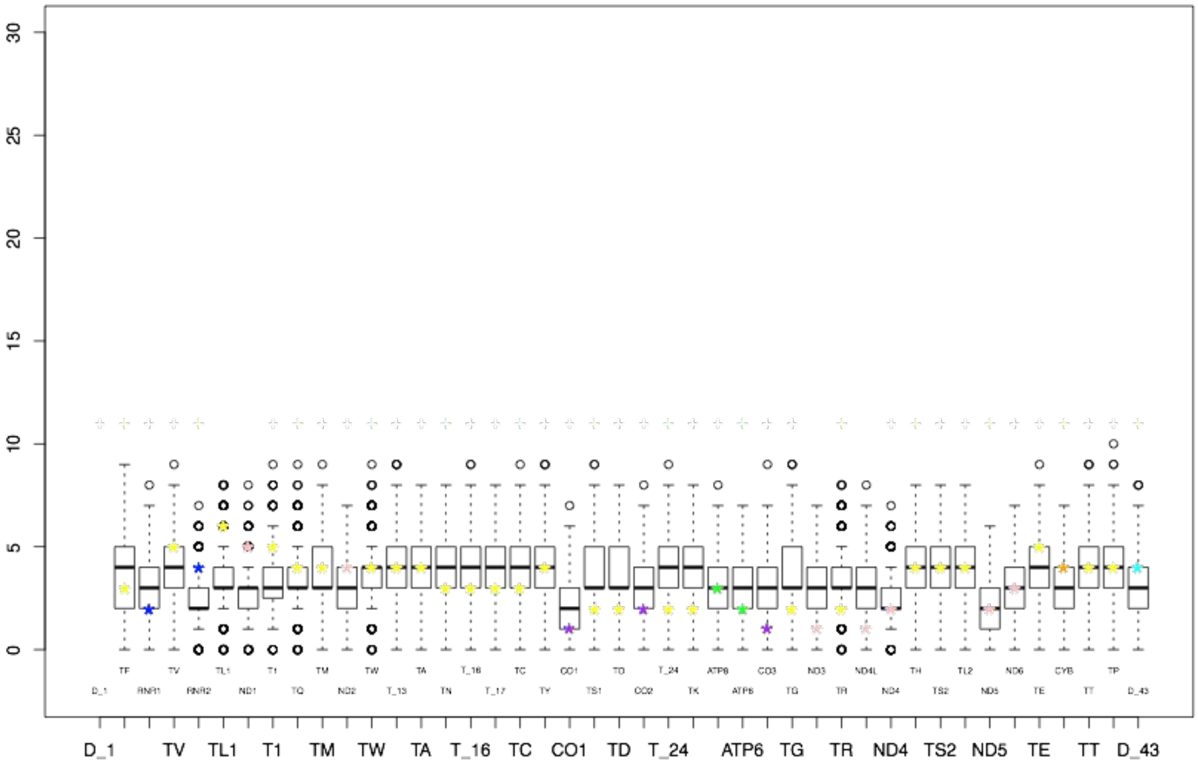


**Fig S8. Enrichment analysis for complete mitochondrial genes integrated into the bonobo genome, relative to human mtDNA sequence.** Data represented as described in Fig S3 for (A) reference-based and (B) polymorphic NumtS.

A

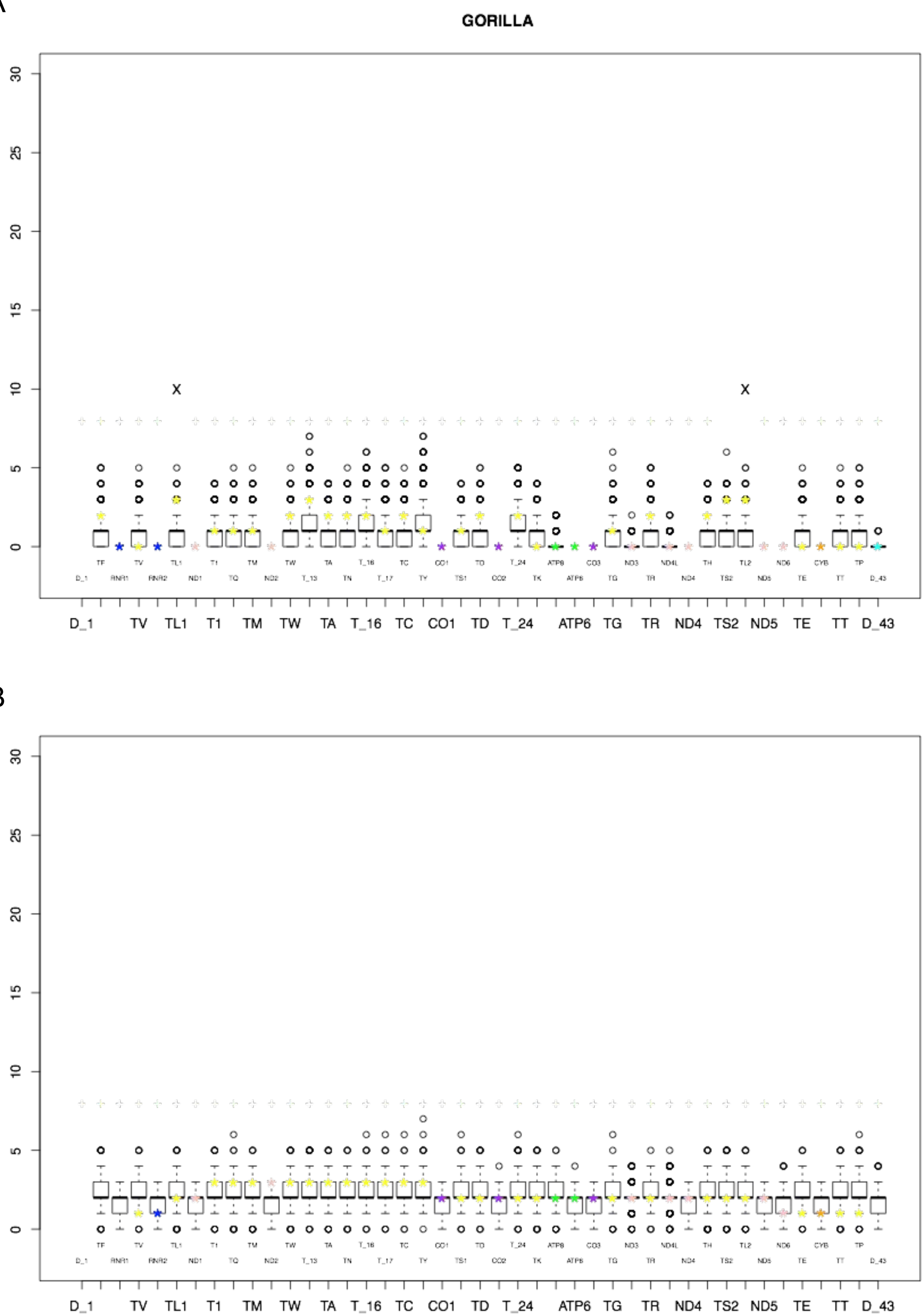


B



**Fig S9. Enrichment analysis for complete mitochondrial genes integrated into the gorilla genome, relative to human mtDNA sequence.** Data represented as described in Fig S3 for (A) reference-based and (B) polymorphic NumtS.

A

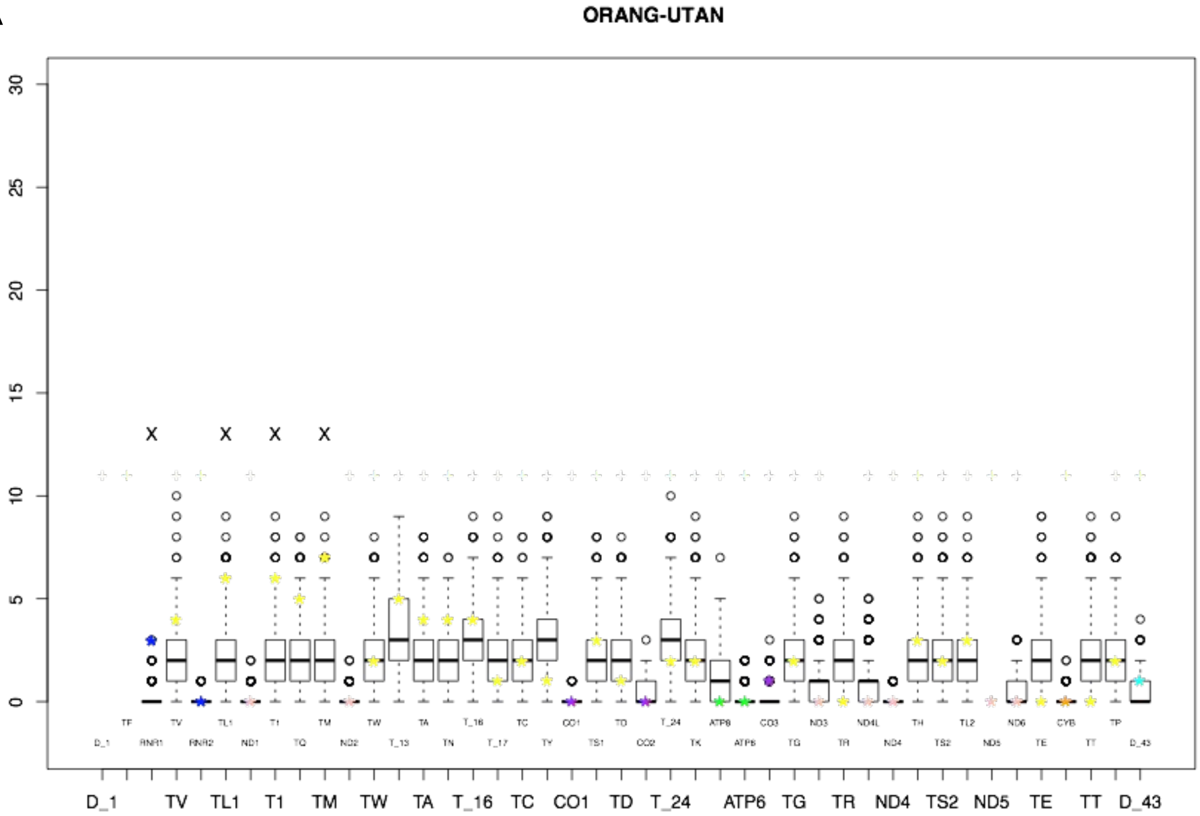


B

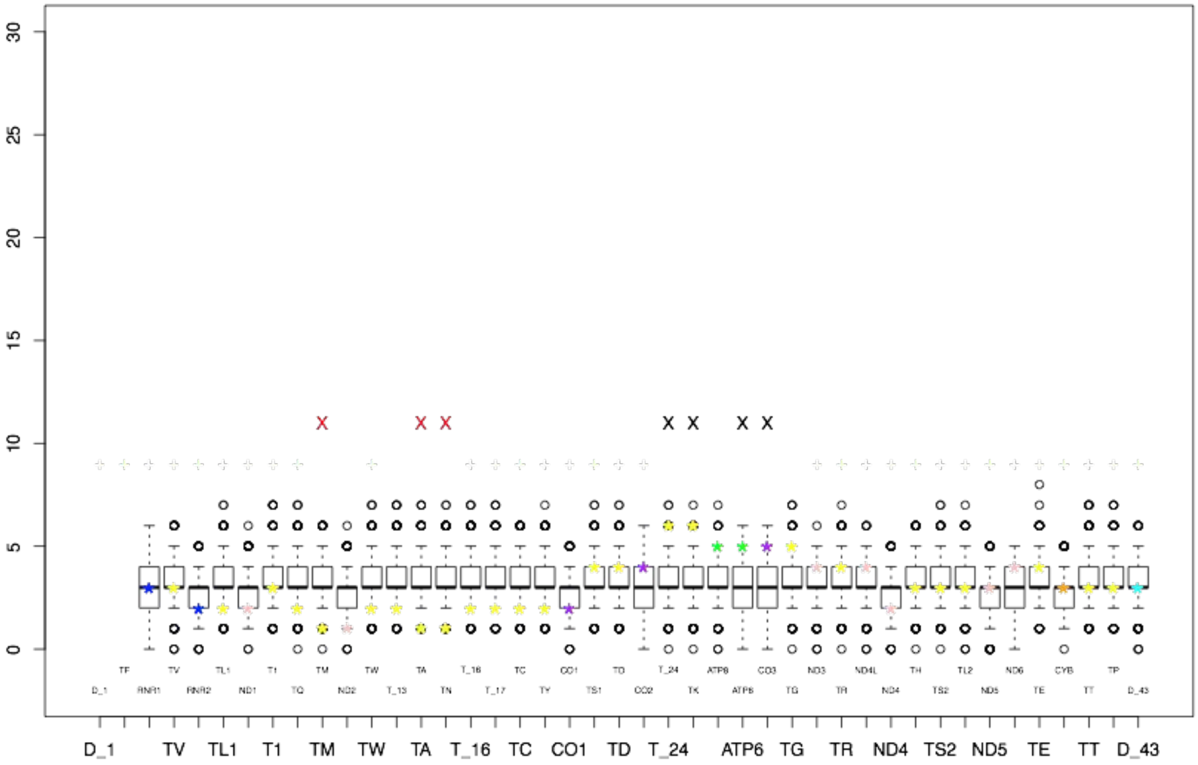


**Fig S10. Enrichment analysis for complete mitochondrial genes integrated into the orang-utan genome, relative to human mtDNA sequence.** Data represented as described in Fig S3 for (A) reference-based and (B) polymorphic NumtS.

A

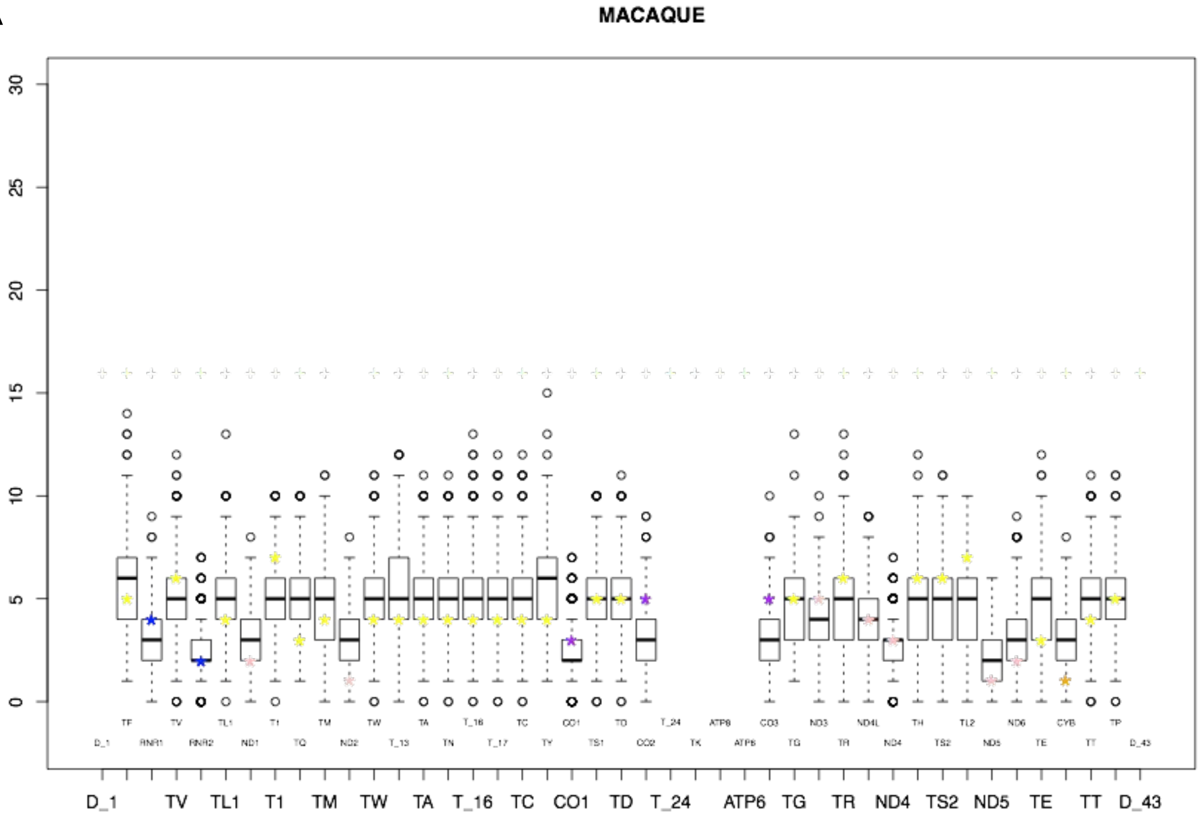


B

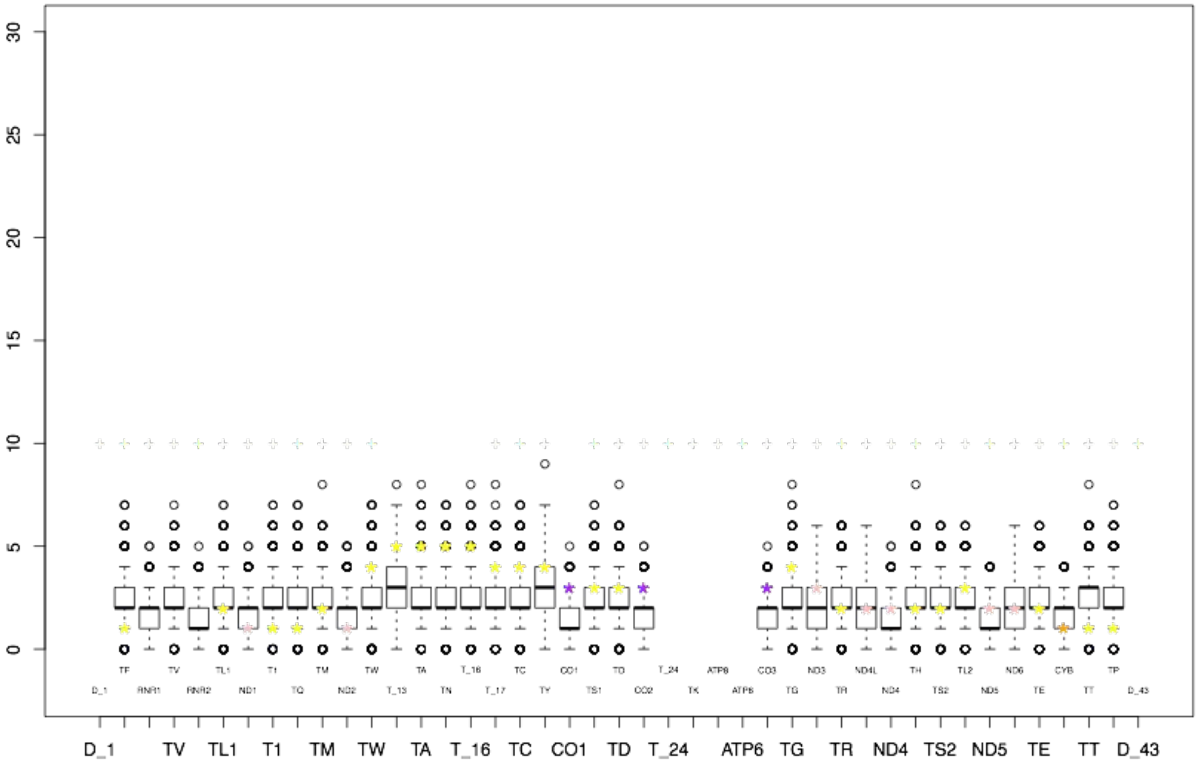


**Fig S11. Enrichment analysis for complete mitochondrial genes integrated into the macaque genome, relative to human mtDNA sequence.** Data represented as described in Fig S3 for (A) reference-based and (B) polymorphic NumtS.
